# Genome plasticity in Papillomaviruses and *de novo* emergence of *E5* oncogenes

**DOI:** 10.1101/337477

**Authors:** Anouk Willemsen, Marta Félez-Sánchez, Ignacio G. Bravo

## Abstract

The clinical presentations of papillomavirus (PV) infections come in many different flavors. While most PVs are part of a healthy skin microbiota and are not associated to physical lesions, other PVs cause benign lesions, and only a handful of PVs are associated to malignant transformations linked to the specific activities of the *E5*, *E6* and *E7* oncogenes. The functions and origin of *E5* remain to be elucidated. These *E5* ORFs are present in the genomes of a few polyphyletic PV lineages, located between the early and the late viral gene cassettes. We have computationally assessed whether these *E5* ORFs have a common origin and whether they display the properties of a genuine gene. Our results suggest that during the evolution of *Papillomaviridae*, at least four events lead to the presence of a long non-coding DNA stretch between the *E2* and the *L2* genes. In three of these events, the novel regions evolved coding capacity, becoming the extant *E5* ORFs. We then focused on the evolution of the *E5* genes in *AlphaPVs* infecting humans. The sharp match between the type of E5 protein encoded in *AlphaPVs* and the infection phenotype (cutaneous warts, genital warts or anogenital cancers) supports the role of E5 in the differential oncogenic potential of these PVs. In our analyses, the best-supported scenario is that the five types of extant E5 proteins within the *AlphaPV* genomes may not have a common ancestor. However, the chemical similarities between E5s regarding amino acid composition prevent us from confidently rejecting the model of a common origin. Our evolutionary interpretation is that an originally non-coding region entered the genome of the ancestral *AlphaPVs*. This genetic novelty allowed to explore novel transcription potential, triggering an adaptive radiation that yielded three main viral lineages encoding for different E5 proteins, and that display distinct infection phenotypes. Overall, our results provide an evolutionary scenario for the *de novo* emergence of viral genes and illustrate the impact of such genotypic novelty in the phenotypic diversity of the viral infections.

## Introduction

Papillomaviruses (PVs) constitute a numerous family of small, non-encapsulated viruses infecting virtually all mammals, and possibly amniotes and bony fishes. According to the International Committee on Taxonomy of Viruses (ICTV: https://talk.ictvonline.org/taxonomy/), the *Papillomaviridae* family currently consists of 53 genera, which can be organized into a few crown groups according to their phylogenetic relationships [31] The PV genome consists of a double stranded circular DNA genome, roughly organized into three parts: an early region coding for six open reading frames (ORFs: *E1*, *E2*, *E4*, *E5*, *E6* and *E7*) involved in multiple functions including viral replication and cell transformation; a late region coding for structural proteins (L1 and L2); and a non-coding regulatory region (URR) that contains the *cis*-elements necessary for replication and transcription of the viral genome. The major oncoproteins encoded by PVs are E6 and E7, which have been extensively studied [39, 40, 65]. However, there is also a minor oncoprotein termed E5, whose functions and origin remain to be fully elucidated [20].

The *E5* ORFs are located in the intergenic region between the *E2* and the *L2* genes. This inter-E2–L2 region is highly variable between PV genomes. In most PV lineages the early and late gene cassettes are located in direct apposition. In a few, non-monophyletic PV lineages, this region accommodates both coding and non-coding genomic segments, which may have gained access to the PV genomes through recombination events with hitherto non-identified donors [7]. PVs within the *Alpha*- and *DeltaPV* genera encode different E5 proteins in the inter-E2–L2 region [6]. Additionally members of the Lambda-MuPV and Beta-XiPV crown groups present in the inter-E2-L2 region large non-coding stretches of unknown significance and/or function [28].

The largest wealth of scientific literature about PVs deals with *AlphaPVs*. These are a clinically important group of PVs that infect primates, and are associated to largely different clinical manifestations: non-oncogenic PVs causing anogenital warts, oncogenic and non-oncogenic PVs causing mucosal lesions, and non-oncogenic PVs causing cutaneous warts. The E5 proteins in *AlphaPVs* can be classified into four different groups according to their hydrophobic profiles and phylogeny [6]. The presence of a given E5 type sharply correlates with the clinical presentation of the corresponding PV infection: viruses that contain E5*α* (*e.g.* HPV16) are associated with malignant mucosal lesions such as cervical cancer; viruses coding for E5*β* (*e.g.* HPV2) are associated with benign cutaneous lesions, commonly warts on fingers and face; and viruses that contain two putative E5 proteins, termed E5*γ* and E5*δ* (*e.g.* HPV6) are associated with benign mucosal lesions such as anogenital warts [6]. Two additional putative E5 proteins, E5∊ and E5*ζ* (PaVE; https://pave.niaid.nih.gov), have been identified in *AlphaPVs* infecting *Cercopithecinae* (macaques and baboons). Contrary to the other E5 proteins, the E5∊ and E5*ζ* are not associated with a specific clinical presentation, although our knowledge about the epidemiology of the infections in primates other than humans is still very limited. It has been suggested that the integration of an *E5* proto-oncogene in the ancestor of (*AlphaPVs*) supplied the viruses with genotypic novelty, which triggered an adaptive radiation through exploration of phenotypic space, and eventually generated the extant three clades of PVs [7].

The only feature that all E5 proteins have in common is their highly hydrophobic nature and their location in the inter-E2–L2 region of the PV genome. It remains unclear whether all E5 proteins are evolutionary related. The E5 proteins of HPV16 and of BPV1 are the only E5s for which the biology is partially known. Despite the absence of sequence similarity, and the differences in immediate interaction partners, the cellular roles during infection are comparable. HPV16 E5 is a membrane protein that localizes in the Golgi apparatus and in the early endosomes. It has been associated to different oncogenic mechanisms related to the induction of cell replication through manipulation of the epidermal growth receptor response [15, 45, 58], as well as to immune evasion by modifying the membrane chemistry [62, 60] and decreasing the presentation of viral epitopes [3]. BPV1 E5 is a very short protein (half the size of HPV16 E5) that also localizes in the membranes. It displays a strong transforming activity, largely by activating the platelet-derived growth factor receptor [19, 44], and it downregulates as well the presentation of viral epitopes in the context of the MHC-I molecules [4].

In this study, we have explored the evolutionary history of the *E5* ORFs found within the inter-E2–L2 region in PVs. First, we identified the PV clades that contain a long intergenic region between *E2* and *L2*, and therewith putative *E5* ORFs. Then, we assessed whether the E5 ORFs in the identified clades originated from a single common ancestor. Next, we verified whether the evolutionary history of the inter-E2–L2 region and of the *E5* ORFs therein encoded is similar to that of the other PVs genes, by comparing their sequences and phylogenies. Finally, we examined whether the different *E5* ORFs exhibited the characteristics of a *bona fide* gene to exclude the conjecture that these are simply spurious translations.

## Methods

### DNA and Protein Sequences

We collected 354 full length PV genomes from the PaVE (pave.niaid.nih.gov) and GenBank (https://www.ncbi.nlm.nih.gov/genbank/) databases (table S1). The corresponding E5 sequences were retrieved from these genomes as well as the intergenic region between the *E2* and *L2* genes (inter-E2–L2). Based on the size of the inter-E2–L2 region in which E5s are present, we selected those with a minimum lenght of 250 nucleotides (fig.1 and fig. S1). For comparison in the tree figures, we extended our analysis and also indicated inter-E2–L2 regions with a minimum length of 125 nucleotides. The URR, *E6*, *E7*, *E1*, *E2*, *L2* and *L1* were also extracted from the collected genomes and analyzed in parallel to the *E5* sequences. We excluded the *E4* ORFs from our analyses as most of its coding sequence overlaps the *E2* gene in a different reading frame and it is supposed to be under different evolutionary pressures [25, 33]. Genes were aligned individually at the amino acid level using MAFFT v.7.271 [34], corrected manually, and backtranslated to nucleotides using PAL2NAL v.14 [61] The alignment was filtered using Gblocks v.0.91b [12]. The URR and the inter-E2–L2 region (non-coding regions) were aligned at the nucleotide level.

### Phylogenetic Analyses

For tree construction of the concatenated *E1*, *E2*, *L2* and *L1* genes, the previously identified recombinant PVs isolated from Cetaceans (PphPV1-2, TtPV1-7, DdPV1, PsPV1) [31, 48, 51] were removed before alignment, leaving us with a data set of 343 PVs. The concatenated E1-E2-L2-L1 alignment was used to construct Maximum Likelihood (ML) trees with RAxML [57] under the GTR+Γ4 model for the nucleotide alignment (fig. S1), using 12 partitions (three for each gene corresponding to each codon position), or under the LG+I+Γ model for the amino acid alignment (fig. 1) using 4 partitions (one for each gene), and using 1000 bootstrap replicates.

**Figure 1.**
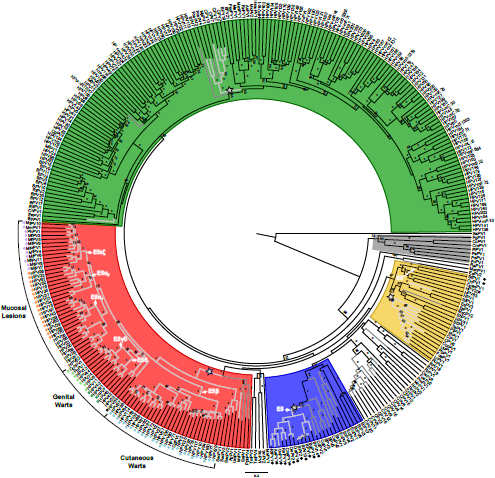
PV phylogenetic reconstruction and identification of clades with an intergenic E2–L2 region. Best-known maximum likelihood phylogenetic tree of the concatenated *E1E2L2L1* amino acid sequences of 343 PVs. Color code highlights the four PV crown groups: red, Alpha-OmikronPVs; green, Beta-XiPVs; yellow, Lambda-MuPVs; blue, Delta-ZetaPVs; gray, a yet unclassified crown group consisting of PVs infecting birds and turtles; and white, PVs without well-supported phylogenetic relationships. Outer labels, *Mucosal Lesions*, *Genital Warts* and *Cutaneous Warts*, indicate the most common tropism for the *AlphaPVs*. Values on branches correspond to ML bootstrap support values. Asterisks indicate a maximal support of 100, and values under 50 are not shown. Branches in light-gray correspond to PV genomes containing an inter-E2–L2 region longer than 250 nt; branches in dark-gray correspond to PV genomes with an inter-E2–L2 region longer than 125 nt. The basal nodes of the four clades containing a relatively long intergenic region between the E2 and the L2 ORFs are labelled with a star. The basal node of the lineages containing an E5 coding sequence is indicated with an arrow, and the corresponding terminal taxa are labelled with a color-coded dot indicating the E5 type. Purple dots indicate: E5*∊ζ*, orange dots: E5*α*, light green dots: E5*γδ*, dark green dots: E5*δ*, blue dots: E5*β*, and black dots are lineages containing unclassified E5 types.

To measure the distances between the URR, *E6*, *E7*, *E1*, *E2*, *E5*, inter-E2–L2, *L2* and *L1* trees, we reduced the data set to 69 PVs so that the taxa are in common among all trees. We reconstructed a phylogenetic tree for each gene separately, as well as for the URR and the inter-E2–L2 region. ML trees were constructed at the nucleotide level using RAxML v.8.2.9 under the GTR+Γ4 model. The weighted and unweighted Robinson-Foulds (RF) distances between trees were calculated [50]. The unweighted RF distance only depends on the topology of the trees, while the weighted RF distance considers edge weights. A correspondence analysis was performed to identify similarities between the topologies of the trees reconstructed for each gene.

### Testing for Common Ancestry using BAli-Phy

In order to evaluate the common ancestry of the E5 ORFs, we used the BAli-Phy algorithm [59]. Under this bayesian framework, the input data are the unaligned sequences, as the alignment itself is one of the parameters of the model to be treated as an unknown random variable [49]. We ran our analysis under the null hypothesis of common ancestry of the intergenic regions. We used the marginal likelihood calculated as the harmonic mean of the sample likelihood to estimate the Bayes Factor between the null hypothesis *Common Ancestry* (CA) and the alternative hypothesis *Independent Origin* (IO) [18]. Therefore, we have ΔBF = log[Prob(CA)]-log[Prob(IO)], such that positive values support CA and negative values support IO. The likelihood for the CA model was obtained running the software for all the E5 sequences together. For the IO scenarios, we ran one analysis for each group independently. We started with the different PV clades that contain an E5 ORF in the inter-E2–L2 region, located within the Alpha-Omikron (red) and Delta-Zeta (blue) crown groups (fig. 1 and fig. S1). In the cases where two putative E5 ORFs were located in the same inter-E2–L2 fragment (for instance for E5*γ* and E5*δ*, and E5∊ and E5*ζ*) sequences were concatenated. Then we ran the analyses on the E5 ORFs within *AlphaPVs* stratifying by the different E5 types that are associated to three distinct clinical presentations; mucosal lesions, cutaneous warts, and genital warts. The values for the independent groups of E5*α*1, E5*α*2, E5*β*, E5*γδ*, E5*δ*, and E5*∊ζ*, and the sum of combinations of these, rendered the likelihood for the IO models. For instance, (*α*1-*α*2-*∊ζ*) + (*γδ*-*δ*) + *β* denotes a hypothesis of three independent ancestries, one tree for the E5 types associated to mucosal lesions (E5*α*1, E5*α*2, and E5*∊ζ* together), another separate tree for the E5 types associated to genital warts (E5*γδ* and E5*δ* together), and another tree for the E5 type associated to cutaneous warts (E5*β*). The likelihood of this example was obtained running BAli-Phy three times: one run for E5*α*1, E5*α*2, and E5*∊ζ*, one for E5*γδ* and E5*δ*, and one for E5*β*. The sum of these three analyses corresponded to the likelihood of the model. We only considered the IO scenarios that were biologically plausible based on the phylogeny of PVs (fig. 1 and fig. S1). The same procedure was applied to the E5 sequences belonging to both the Alpha-Omikron and Delta-Zeta crown groups. This analysis was performed at the amino acid level using the LG substitution model. For each model, three independent MCMC chains were run for at least 100000 iterations. The three runs were combined and checked for convergence.

### Random permutations to test for Common Ancestry

To support the results of the BAli-Phy analyses, we performed a random permutation test as described in de Oliveira Martins and Posada 2016 [17]. In this test the sequences for one of the groups are randomly shuffled and statistics are recalculated after realignment with MUSCLE [24], which tells us how much the results using the original data departs from those with phylogenetic structure partially removed. The statistics used in this test are ML tree length and Log Likelihood calculated with PhyMLv3.0 [32]. As for the BAli-Phy test, these analyses were performed at the amino acid level using the LG substitution model. We obtained a distribution by reshuffling one of the groups (for example the E5*∊ζ* sequences) 100 times, each time realigning against the other groups from the data set, and comparing the resulting phylogeny with those if we separate again the groups. For each iteration, the alignment is always optimised and the statistics are calculated. To make the statistics comparable, the same alignment is used for both the IO and CA hypotheses. We compare the distribution for the CA and IO hypotheses with a Kruskal-Wallis rank sum test and a multiple comparison test after Kruskal-Wallis. The results were confirmed by performing Wilcoxon rank sum tests with continuity correction. Lower ML tree length and superior Log likelihood values are expected to support the best model.

### Generation of Random ORFs

In order to assess whether the *E5* sequences were larger than expected by chance, we estimated first the median A/T/G/C composition of the inter-E2–L2 regions of *AlphaPVs* (A:0.22; T:0.41; G:0.20; C:0.17). Using in-house perl scripts, we created a set of 10,000 random DNA sequences with this median nucleotide composition and with a median length of 400 nt. Then, we computed the length of all putative ORFs that may have appeared in this set of randomly generated DNA sequences.

### dN/dS Values

To determine whether the *E5* ORFs are protein-coding sequences, we computed the dN/dS values for all *E5* ORFs as well as for the other PV ORFs (*E1*, *E2*, *E6*, *E7*, *L1*, *L2*). The dN/dS values were computed with SELECTON (http://selecton.tau.ac.il/overview.html [23], using the MEC model [22]. The likelihood of MEC model was tested against the M8a model [71], which does not allow for positive selection. As these models are not nested, AIC scores were compared. For all the sequence sets, the MEC model was preferred over the M8a model.

### Pairwise Distances

To evaluate the diversity of the *AlphaPV* genes, we calculated the pair-wise distances between aligned sequences within each group of the *E5* ORFs, the other PV ORFs (*E1*, *E2*, *E6*, *E7*, *L1*, *L2*), and the URR. These random intergenic CDS were generated by extracting the non-coding region of the E2–L2 fragments of all *AlphaPVs*. Then, for each non-coding region, we extracted a random subregion with the same length as the *E5* ORF of this PV. These random intergenic regions were truncated at the 5’ to get a sequence length multiple of 3. All internal stop codons were replaced by N’s. Pair-wise distances between aligned DNA sequences were calculated using the TN93 model. All distances were normalized with respect to the corresponding one obtained for *L1*.

### Codon Usage Preferences

We calculated the codon usage preferences (CUPrefs) for the *E5 AlphaPV* ORFs. The frequencies for the 59 codons with redundancy (i.e. excluding Met, Trp and stop codons) was retrieved using an in-house perl script. For each of the 18 families of synonymous codons, we calculated the relative frequencies of each codon. We performed the same analysis for all other ORFs in the same genomes (*E1*, *E2*, *E6*, *E7*, *L1* and *L2*) as well as to the randomly generated intergenic CDS. A matrix was created in which the rows corresponded to the ORFs on one PV genome and the columns to the 59 relative frequency values, such that each row had the codon usage information for a specific ORF. We performed a non-metric Multidimensional Scaling (MDS) analysis with Z-transformation of the variables in order to assess similarities in codon usage preferences of the *E5* ORFs with respect to the other *AlphaPV* ORFs, as described in [25]. In parallel, we performed a two-step cluster analysis with the same relative frequency values. The optimal number of clusters was automatically determined using the Bayesian Information Criterion (BIC).

### GRAVY Index

For all E5 proteins the grand average hydropathy (GRAVY) was calculated by adding the hydropathy value for each residue and dividing this value was by the length of the protein sequence [36].

### Statistics and Graphics

Statistical analyses and graphics were done using R [46], with the aid of the packages “ape”, “ade4”, and “phangorn”. The final display of the graphics was designed using Inkscape v.0.92 (https://inkscape.org/en/).

## Results

### Do the E5 ORFs Present in the Genomes of PVs Belonging to Different Crown Groups Have a Common Ancestor?

We collected 354 full length PV genomes from the PaVE (pave.niaid.nih.gov) and GenBank (https://www.ncbi.nlm.nih.gov/genbank/) databases (table S1). After removing eleven recombinant sequences we constructed a maximum likelihood phylogenetic tree of the concatenated *E1E2L2L1* sequences at the nucleotide and amino acid levels. Out of the 354 PV genomes, we identified 339 with an intergenic region (of at least 1 nucleotide) between the *E2* and *L2* genes. Of these, 83 contain an E5 ORF in the inter-E2–L2 region (fig. 1 and fig. S1). The E5 ORFs have a median size of 144 nucleotides (min: 126, max: 306). Based on the size of inter-E2–L2 region in which E5s are present (min: 289, median: 517, max: 938), we identified four PV clades containing an intergenic region selecting for a minimum size of 250 (min: 262, median: 512, max: 1579). This threshold is below the minimum of 289 nucleotides to allow for inclusion of possible unidentified PV lineages containing unknown E5-like ORFs in the inter-E2–L2 region. The identified clades are indicated with a star in fig. 1 and fig. S1, and are located in the four PV crown groups: Alpha-Omikron (coloured red), Delta-Zeta (coloured blue), Lambda-Mu (coloured yellow), and Beta-Xi (coloured green). Additionally, three recombinant bottlenose dolphin PVs (TtPV1-3) belonging to the *UpsilonPV* genus, also present an inter-E2–L2 region. Only the clades identified in the Alpha-Omikron and Delta-Zeta crown groups, have an E5 ORF present within the inter-E2–L2 region. The two other clades that locate within the Lambda-Mu and Beta-Xi crown groups also contain this relatively long intergenic region. Although, for these clades the inter-E2–L2 region does not contain any apparent ORFs. Interestingly, an ORF named *E5* is present in the Lambda-Mu clade in two rabbit PV genomes (SfPV1 and OcPV1), where no intergenic non-coding region is present and E5 largely overlaps with both the *E2* and *L2* genes in the case of SfPV1 and with *L2* in the case of OcPV1. There are other cases, like HPV16, where E5 partially overlaps with the *E2* gene. Nonetheless this overlap is small (4 nucleotides) compared to the almost complete overlap of E5 ORFs with *L2* in the rabbit PV genomes. All things being equal, the E5 ORFs in the rabbit PV genomes seem unique in a way that no inter-E2–L2 region is present at all.

In order to determine whether the E5 ORFs in the different PV crown groups share a single common ancestor, we tested for common ancestry using BAli-Phy as described in de Oliveira Martins and Posada 2014 [18]. We named the clades according to their coloured crown groups, therefore we have the red clade (including 69 E5 sequences), the blue clade (12 E5 sequences), and the yellow clade (2 E5 sequences). For the common ancestry test, trees are inferred for all groups combined as well as separately (see Materials an Methods). Therefore, we could not include the yellow clade in this test, as this clade contains only two sequences and no trees can be inferred. We performed the analysis on the full data set (excluding the two yellow clade sequences) containing 81 sequences and on a reduced dat set containing 24 sequences; twelve representative E5 sequences from the red clade and the twelve E5 sequences from the blue clade. We made the choice between the alternative hypotheses *Common Ancestry* (CA) and *Independent Origin* (IO) by computing the marginal likelihoods using the stabilized harmonic mean estimator. We ran our analysis under the null hypothesis of CA of the E5 ORF. Therefore, we have ΔBF = log[Prob(CA)] - log[Prob(IO)], such that positive values support CA and negative values support IO. Other statistics that we take into account are the alignment length and the Bayesian tree length, calculated as the sum of the branch lengths. For both the alignment length and tree length, lower values support the best model.

The results are contradictory between the different statistics tested. On the one hand, based on the likelihood the best supported model is CA for the E5 ORFs in the Alpha-Omikron and Delta-Zeta PV crown groups (table 1). Nonetheless, the difference in Log likelihood (ΔBF) between the CA and IO hypotheses is very small for both the full and reduced data sets. On the other hand, the alignment length and tree length statistics support the IO hypothesis. Previous approaches of other type of CA tests have shown to give misleading conclusions on alignments without any phylogenetic structure [35], as well as on unrelated families of protein coding sequences [72]. As these approaches all started from a fixed alignment there could be an initial bias towards CA [18, 63, 73]. The BAli-Phy approach used here partly reduces this bias, as it starts from unaligned sequences and estimates simultaneously the alignment and the phylogeny. Given the inconclusive results, we performed a random permutation test as described in de Oliveira Martins and Posada 2016 [17]. In this test the columns of the alignment for one of the groups are randomly shuffled and statistics are recalculated after realignment. Contrary to the BAli-Phy test, all trees are produced within a maximum likelihood (ML) framework (see Materials and Methods). We performed this test on both the full and the reduced data sets, using 100 iterations. For each iteration we recovered the ML tree length and Log likelihood, and estimated the empirical distribution of these value. If the E5 ORFs have an IO, we expect lower ML tree length and superior Log likelihood values for this hypothesis (H1). Our results show that for both the full and reduced data sets we obtained significant differences between the ML tree length distributions of CA and IO, where the IO hypothesis is favoured fig. S2A-B. However, for the Log likelihood distributions there is no significant difference between CA and IO for the full data set, and for the reduced data set the CA hypothesis is slightly favoured fig. S2C-D.

**Table 1.** Hypothesis testing on the origin of the E5 ORFs in the Alpha-Omikron (red) and Delta-Zeta (blue) PV crown groups. For each hypothesis tested, common ancestry (H0) and independent origin (H1), we show the marginal likelihood (P(data|M)) value, the ΔBF, the alignment (ali) length and tree length. Cells highlighted in gray indicate the best-supported scenario for the respective statistic.

|  | full data set |  |  |  | reduced data set |  |  |  |
| --- | --- | --- | --- | --- | --- | --- | --- | --- |
| Model | P(data\|M) | ΔBF | ali length | tree length | P(data\|M) | ΔBF | ali length | tree length |
| H0: (red-blue) | -7129.167 | 0 | 402 | 26.946 | -2608.666 | 0 | 344 | 10.893 |
| H1: red + blue | -7139.029 | 9.862 | 373 | 24.301 | -2617.114 | 8.448 | 335 | 10.706 |

The initial idea of the permutation test was to resort only to simple summary statistics such as the ML tree length, rather than to rely on Log likelihood values [17]. If we only regard the ML tree length values, IO for the red and blue clades is suggested to be the best supported model. Nevertheless, we can not ignore the Log likelihood values of the permutation test nor the results of the BAli-Phy test, and therefore, we cannot make a conclusive choice between the alternative hypotheses CA and IO. Finally, when we look at the final trees produced by BAli-Phy fig. S3, we observe that the branch lengths leading to each group are long compared to the other branches, suggesting that IO is the preferred model. In addition, these trees suggest that the E5 ORFs within the *AlphaPV* (red clade) do not originate from a single CA, which may have introduced a bias in our CA test.

### Do the E5 ORFs Present in the Genomes within the AlphaPV Clade Have a Common Ancestor?

In the *AlphaPV* clade within the Alpha-Omikron crown group (red), the six E5 types are present in five different clades (fig. 1 and fig. S1). E5*α* exists in two different clades of PVs associated to mucosal lesions, hereafter named E5*α*1 and E5*α*2, consisting of eight and nine sequences respectively. E5*β* is present in all PVs associated to cutaneuous warts, consisting of 28 sequences. E5*δ* exists in all PVs associated to anogenital warts. Of these, only four PV genomes contain E5*δ* in isolation. The other seven PV genomes contain two E5 types; E5*γ* and E5*δ*, hereafter named E5*γδ*. Finally, E5*∊ζ* is present in twelve non-human *AlphaPV* genomes that infect *Cercopithecinae* and that are associated to mucosal lesions.

The BAli-Phy trees obtained in the CA test above, suggest that the E5 ORFs within *AlphaPVs* may have an IO (fig. S3). These trees, that are based on the E5 amino acid sequences, show a clear separation depending on the clinical presentation of the infections: mucosal lesions (E5*α*1, E5*α*2, and E5*∊ζ*), genital warts (E5*γ*, and E5*γδ*), and cutaneous warts (E5*β*). One exception is HPV54, which has an unclassified E5 type and is associated to genital warts. This PV clusters with the E5*α*1 type of mucosal lesions. To address whether the E5 ORFs present in the genomes of the *AlphaPVs* have a CA, we applied the same procedures as described above. We considered different plausible IO scenarios based on the E5 types and the phylogeny of the *AlphaPVs* (fig. 1 and fig. S1). The BAli-Phy analysis showed that the CA hypothesis was the best-supported model for all statistics, while the hypothesis of each clade having an IO (H6) had the lowest support (table 2). The second best-supported IO model H1 -where E5*β* has an IO– has a small difference in Log likelihood with the CA model (H0). As in the results described above, the random permutation tests disagree with the results of the BAli-Phy approach. The results of the random permutation test suggest that based on ML tree lenght the IO H6 is the best supported model, while based on Log likelihood the CA model (H0) and IO H1 model are equally probable (fig. S4). Although the CA tests performed here give inconclusive results, the IO H1 model is also supported by the trees produced, where long branches separate E5*β* and the other E5 types. In this scenario E5*α*1, E5*α*2, *γδ*, E5*δ*, and E5*∊ζ* (encoded in PVs with mucosal and anogenital tropism) have a CA, but E5*β* (encoded in PVs with cutaneous tropism) has an IO. We therefore propose that at least E5*α*1, E5*α*2, *γδ*, E5*δ*, and E5*∊ζ* have a single ancestor, and originated from the same recombination donor and/or gained access to the ancestral genome through a single integration event. Further tests are needed to conclude whether E5*β* originated from the same ancestor as the other E5 types or whether it has an independent origin.

**Table 2.**
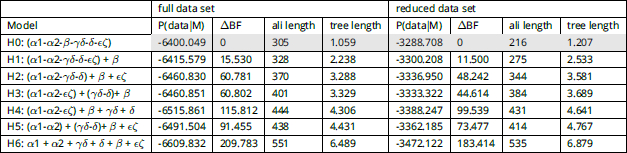
Hypothesis testing on the origin of the E5 ORFs within the AlphaPV clade (red). For each hypothesis tested, common ancestry (H0) and independent origins (H1-H6), we show the marginal likelihood (P(data|M)) value, the ΔBF, the alignment (ali) length and tree length. Cells highlighted in gray indicate the best-supported scenario for the respective statistic.

### In *AlphaPVs*, The Evolutionary History of The inter-E2–L2 Region is Different from That of E5

In order to look deeper into the evolutionary history of the inter-E2–L2 region within *AlphaPVs*, we performed phylogenetic analyses and compared the tree topology for the inter-E2–L2 fragment sequences and the *E5* ORF with the topologies obtained for each of the PV ORFs (*E6*, *E7*, *E1*, *E2*, *L2* and *L1*) as well as for the non-coding URR. We calculated the weighted and unweighted Robinson-Foulds (RF) distances between paired trees and we performed a correspondence analysis in order to identify similarities among the topologies of the PV gene trees (fig. 2). The first axis captured a large fraction of the variance (more than 50% in the weighted RF distance) and splitted the *E5* and the inter-E2–L2 reconstructions from those of core PV genes. The second axis contained more than 15% of the overall variance and splitted the topologies of the early genes *E6*, *E7*, *E1* and *E2*, from those of the late genes *L2* and *L1*, and the URR. Interestingly, in this second axis the inter-E2–L2 clustered together with the late genes, while the *E5* genes clustered together with the early genes. These results suggest that the inter-E2–L2 region and E5 may have different evolutionary histories.

**Figure 2.**
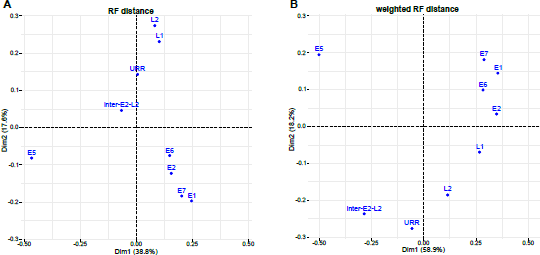
Correspondence analysis of the unweighted (A) and weighted (B) Robinson-Foulds tree distance comparing maximum likelihood trees construced for each of the PV ORFs, the inter-E2–L2 region, and the URR.

### The E5 ORFs in *AlphaPVs* Display the Characteristics of a Genuine Gene

Since it is often discussed whether the *E5* ORFs in *AlphaPVs* are actual coding sequences, we performed a number of analyses in order to assess whether the different *E5* ORFs exhibit the characteristics of a *bona fide* gene. To determine whether the *E5* ORFs are larger than expected by chance, we constructed first 1000 random DNA sequences with the same median nucleotide composition as the inter-E2–L2 region of *AlphaPVs*, we identified all putative ORFs appearing by chance in these randomly generated DNA sequences and we computed their nucleotide length. (fig. 3) shows the cumulative frequency of the *E5* genes length and of the random ORFs. A one-way ANOVA followed by a post-hoc Tukey honestly significant difference test was performed, with *gene* as a factor (table S2) shows that ORFs in randomly generated sequences are shorter than any of the E5 ORFs (Tukey HSD: *p* < 0.0001).

**Figure 3.**
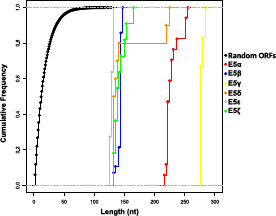
Cumulative frequency of the nucleotide length for each group of the *E5* genes and random ORFs. The different types of E5 are colour-coded as indicated in the legend.

Besides length, evidence of selective pressure is another signature of *bona fide* genes. We calculated the dN/dS values for all *E5* sequences (fig. 4). Our results showed that the *E5* genes display a dN/dS distribution that is significantly lower than 1 (Wilcoxon-Mann-Whitney one side test: p < 0.001), with median values ranging from 0.13 to 0.40. All other PV genes presented median dN/dS values lower than the *E5* sequences (Tukey HSD: *p* < 0.001) (fig. 4).

**Figure 4.**
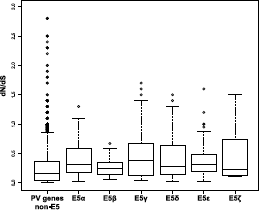
dN/dS values for each group of the *E5* genes and the other PV genes (E1, E2, E6, E7, L1 and L2).

We next calculated the pair-wise distances between terminal taxa for all ORFs and for the URR in *AlphaPVs*, as well as for a set of randomly generated intergenic CDS (fig. 5). These random CDS were generated using the average nucleotide composition from the inter-E2–L2 region of *AlphaPVs*, selecting for the same length distribution as the E5 ORFs (see Materials and Methods). Pairwise distances were normalized with respect to the corresponding *L1* distance. The highest rates of variation were found as expected in the random intergenic CDS region and the lowest rates in the PV genes that are not *E5* (Tukey HSD, *p* < 0.001). Our results also showed that all *E5* genes presented lower rates of variation than the random intergenic CDS but higher rates than the other PV genes. The E5*α*, E5*β* and E5*ζ* showed higher rates of variation compared to the URR (Tukey HSD, *p* < 0.001). Contrary, the E5*γ*, E5*δ*, and E5*E* showed lower rates of divergence in comparison to the URR (Tukey HSD, *p* < 0.001).

**Figure 5.**
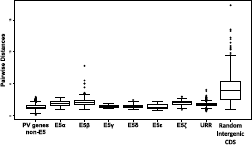
Pairwise distances between *AlphaPVs* for the all genes, the URR, and a set of randomly generated intergenic CDS. All values have been normalized to the corresponding *L1* pairwise distances.

In PVs, codon usage preferences (CUPrefs) are different from those of their hosts, and viral genes with similar expression patterns display similar CUPrefs [25]. To corroborate whether the CUPrefs of the *E5* genes are similar to those of the other PV genes, we calculated the relative frequencies of the 59 codons in synonymous families in the *E5* genes and in the rest of PV genes and the randomly generated intergenic CDS. Then we performed a multidimensional scaling (MDS) analysis on the 59-dimensional codon usage vectors, and in parallel, an unsupervised two-step cluster analysis (fig. 6). The optimal number of clusters was three: one cluster containing the early *E1* and *E2* genes; a second cluster containing late *L2* and *L1* genes; and a third cluster containing the *E5*, *E6*, *E7* oncogenes.

**Figure 6.**
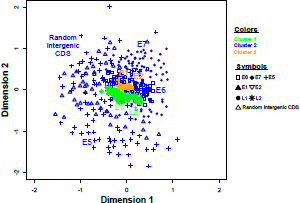
Multidimensional Scaling (MDS) plot of codon usage preferences for the *AlphaPV* ORFs. The ORFs were independently clustered by an unsupervised two-step clustering algorithm. The best assembly included three clusters, displayed onto the MDS plot as with a color code, composed respectively by the oncogenes *E5, E6* and *E7*; the early genes *E1* and *E2*; and the capsid genes *L1* and *L2*.

As the best-studied E5 proteins are transmembrane proteins, we hypothesized that a *bona fide* E5 protein should be more hydrophobic than expected by chance. We calculated the GRAVY index for the E5 proteins as well as for the ORFs enconded in the randomly generated intergenic CDS (fig. 7). We found that E5*α*, E5*β*, E5*γ*, E5*δ*, and E5*E* are more hydrophobic than the random intergenic CDS (Wilcoxon-Mann-Whitney test, *p* < 0.0001). The E5*ζ* is the only E5 protein that did not tested significantly more hydrophobic than the random intergenic CDS (Wilcoxon-Mann-Whitney, *p* = 0.125).

**Figure 7.**
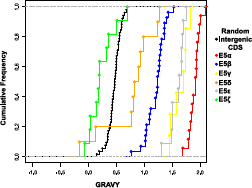
Cumulative frequency of the GRAVY index for the *E5* ORFs and the randomly generated intergenic CDS.

## Discussion

Reconstructing how PV genes have originated and evolved is crucial for explaining the genetic basis of the origin and evolution of phenotypic diversity found in PVs, if we eventually aim to understand why certain PVs are oncogenic while their close relative cause anodyne infections. In this work our first aim was to study the origin of the *E5* oncogenes in *AlphaPVs*. This viral genus hosts around fifty viral genotypes with a relative narrow host distribution (they seem to be restricted to Primates), but with very diverse phenotypic presentations of the infections: many of them are associated to asymptomatic infections of the skin, but also of the oral, nasal, or anogenital mucosas; some of them cause productive infections that result in common skin warts, or in genital warts; and a number of them cause chronic infections that may result in anogenital or oropharyngeal cancers [21, 27]. All *AlphaPVs* present a region between the *E2* and *L2* genes, potentially encoding in all cases for conserved ORFs. With few exceptions [10], actual gene expression and protein function for *E5* oncogenes have only been characterized for the more oncogenic HPVs, which carry E5 proteins of type E5*α* [6]. These E5*α* behave as oncoproteins, promoting cell division and allowing the infected cells to avoid immune recognition [3, 62, 60].

Since all the *E5* ORFs in *AlphaPVs* map between the *E2* and *L2* genes we extended our analysis to the evolution of this intergenic region in the Alpha-Omikron crown group. Finally, since a number of non-monophyletic PVs also contain a sometimes long non-coding region between the *E2* and *L2* genes in their genomes that may also encode for genes named *E5*, we expanded our analyses to the full set of PV sequences containing a long non-coding region at this genomic location. PVs displaying an intergenic region between *E2* and *L2* are not monophyletic, and belong instead to four polyphyletic clades in the PV tree (fig. 1 and fig. S1). It can be argued that the ancestral PV genomes could have already presented an inter-E2–L2 region, which may have undergone several loss events. Such repeated losses have been invoked as a mechanism to explain the repeated absence of early genes (*E6* and *E7*) in certain PVs [66]. Alternatively, the different inter-E2–L2 regions present in extant PV genomes could derive from one or from several genetic events in which an ancestral sequence could have gained access to one ancestral PV genome.

We can formulate two main non-exclusive mechanisms to explain the origin of the four extant groups of inter-E2–L2 regions in the PVs genomes: random nucleotide addition and recombination. Random nucleotide addition is a plausible mechanism, based on the way the PV genome replicates. The replication of the PV genome occurs bidirectionally during the non-productive stages of the infection, yielding episomes [26]. During the PV bidirectional replication, the replication forks start at the URR and converge opposite to the origin of replication, which happens to lay between the *E2* and *L2* genes. At this point, concerted DNA breaks are required for decatenation, which eventually generates two separate circular dsDNA molecules. The end joining of these DNA breaks is error prone. Indeed, the DNA close to the break site can be used as a template for *de novo* synthesis before the DNA ends are joined, resulting in the non-templated introduction of a stretch of additional nucleotides [52], which may lead to the emergence of an ancestral inter-E2–L2 region in one or in several instances during the evolutionary history of PVs.

Recombination can also be invoked as a mechanism that could result in the integration of novel DNA sequences into the PV genome. In parallel to the host keratinocyte differentiation, replication of the viral genome switches from bidirectional to unidirectional [26, 37], generating large linear molecules of concatenated viral genomes [16]. Unidirectional replication relies on homologous recombination, as this mechanisms is required for resolving, excising and recircularizing the concatenated genomes into individual plasmid genomes [29, 38, 54]. Additionally, productive replication concurs with a virus-mediated impairment of the cellular DNA damage repair mechanisms [14, 68], thus rendering the overall viral replication process error-prone by increasing the probability of integrating exogenous DNA during recircularization. Phylogenetic evidence for the existence and fixation of such recombination events is provided by the incongruence in the reconstruction of the evolutionary history for different regions of the PV genome. In all cases, such inconsistencies appear when comparing the phylogenetic inference for the early and for the late genes of the genome, respectively upstream and downstream the recombination-prone genomic region. Evidence for recombination has been described at several nodes in the PV tree. The first example occurs at the root of *AlphaPVs*, with the species containing oncogenic PVs being monophyletic according to the early genes (involved in oncogenesis and genome replication), and paraphyletic according to the late genes (involved in capsid formation) [6, 41]. The second example is provided by certain PVs infecting cetaceans, which display the early genes related to those in other cetacean PVs in the Alpha-Omikron crown group (in red in fig. 1) and the late genes related to those in bovine PVs in the Beta-Xi crown group (in green in fig. 1) [30, 48, 51]. Finally, the most cogent examples of recombination between distant viral sequences are two viruses isolated from bandicoots and displaying the early genes related to Polyomaviruses and the late genes related to PVs [5, 70].

The inter-E2–L2 sequences may occasionally be very long and span more than 1 Kbp, a considerable size for an average genome length of around 8 Kbp. Additionally, for many viral genomes, the sequences in the inter-E2–L2 region do not resemble other sequences in the databases, and do not seem to contain any functional elements, neither ORFs nor transcription factor binding sites or conserved regulatory regions [28, 47, 56]. Despite the lack of obvious function and of their length, these sequences seem to belong *bona fide* in the viral genome in which they are found, as they are fixed and conserved in viral lineages [47]. Although the two hypothesis referred above to explain the origin of the inter-E2–L2 regions (random nucleotide addition and recombination) are plausible, we interpret that the presence of long and conserved sequences in certain monophyletic clades (labeled with a star in fig. 1 and fig. S1) suggests that the respective insertions of each of these long sequences in the ancestral genomes occurred during single episodes, pointing thus towards a recombination event.

The putative ORFs that emerged in the inter-E2–L2 region are often named *E5*. Notwithstanding, our results suggest the E5 proteins encoded in the different clades may not be monophyletic. Specifically, this would imply that the *E5* ORFs in *AlphaPVs* (*e.g.* HPV16 *E5*) are not evolutionarily related to the *E5* ORFs in *DeltaPVs* (*e.g.* BPV1 *E5*). This is an important change in perspective, because these two proteins are often referred to and their cellular activities compared as if they were orthologs [2, 67]. Yet, the E5 sequences are short and display similar amino acid composition because of their transmembrane nature, and these two facts combined reduce the power of the algorithms used to pinpoint common ancestry between genes. Further tests are needed to resolve the riddle on the origin of E5s: either *in silico* by improving the CA test or experimentally by evolving a predicted ancestor(s) of E5 or by performing *de novo* gene evolution on the inter–E2-L2 region.

When restricting our analysis to theE5 ORFs within the *AlphaPVs*, we found support for monophyly (table 2), indicating that a single event on the backbone of the ancestral *AlphaPV* genome could have led to its emergence. Nevertheless, the alternative hypothesis of E5*β* having an independent origin was not significanlty worse. This hypothesis is supported by the different tropism of lineages within *AlphaPVs*: those containing an E5*β* display an essentially cutaneous tropism, while all other lineages encoding for E5*α*, E5*γ*, E5*δ*, E5∊, and E5*ζ*, display a mucosal tropism. Indeed, there is no evident sequence similarity between the E5 proteins, inasmuch as the evolutionary divergence between E5*β* and the other *E5* ORFs rises to 80% [6]. Phylogenetic reconstruction based on the *E5* ORFs showed a star-like pattern with the main branches emerging close to a putative central point [6]. These features could be related to the multiple ancestries of the different *E5* ORFs.

It remains unclear how the different *E5* genes emerged in the *AlphaPV* genomes. Our interpretation based on the evidence here provided is as follows. Under the hypothesis of recombination, within *AlphaPVs*, a non-coding sequence was integrated in a single event between the early and the late genes in the genome of an ancestral PV lineage, which infected the ancestors of Old World monkeys and apes. Mutations in this originally non-coding region gave birth to the different *E5* ORFs. Such *de novo* birth of new protein-coding sequences from non-coding genomic regions is not unfamiliar and has been reported in for example *Drosophila* [42, 74], yeast [8] and mammals [64]. Experimentally, it has been shown that random, E5-like short peptide sequences can indeed insert in the cellular membranes and display a biological activity [13]. Using genetic selection, these small artificial transmembrane amino acid sequences that do not occur in nature were able to bind and activate the platelet derived growth factor (PDGF) *β* receptor (just like BPV E5 does), resulting in cell transformation and tumorogenicity [13]. Therefore we consider *de novo* birth of the *E5* genes in the inter-E2– L2 region a plausible hypothesis. The randomly appeared *E5* genes, short and enriched in hydrophobic amino acids, could thus have provided with a rudimentary function by binding to membrane receptors or by modifying membrane environment. Such activities may have lead to an increase in viral fitness and could have been selected and enhanced, resulting in the different *E5* genes lineages observed today.

The location within the inter-E2–L2 region and the hydrophobic nature of the protein have up to date been the criteria to classify the *E5* ORFs as putative genes. This is probably the reason for which we found all *E5* ORFs, with the only exception of E5*ζ*, more hydrophobic than expected by chance (fig. 7). However, for most of these ORFs we do not have evidence of their expression *in vivo*. Moreover, the possible independent origins of *E5*, raise the concern of whether all *E5* ORFs are actually coding sequences. In this study, we have used several approaches in order to distinguish true *E5* genes from spurious ORFs that are not functional. As orthologs of the *E5* genes are not found in other viruses or in their hosts, we have studied the *E5* ORFs in the context of orphan genes. In agreement with studies of orphan genes in other species [11, 64, 69], the *E5* genes are shorter than the other PV genes. It has previously been proposed that there is a direct relationship between the length of a gene and its age [1, 43, 64]. However, a *bona fide* gene should be longer than expected by chance [55], and this is what we actually find for the different *E5* ORFs (fig. 3).

For a new functional protein to evolve from randomly occurring ORFs, it needs to be produced in significant amounts. These proteins are expected to evolve under neutral selection, as these are unlikely to be functional at first. By combining ribosome profiling RNA sequencing with proteomics and SNP information Ruiz-Orera *et al.* found evidence to support this hypothesis [53]. By analyzing mouse tissue they found hundreds of small proteins that evolve under no purifying selection. Regarding the *E5* ORFs, we obtained dN/dS ratios below 1 (fig. 4), indicating negative or purifying selection, reinforcing the idea that extant E5s may be functionally relevant. Gene CUPrefs have a strong effect on ORF translation, where a favorable codon composition may facilitate the translation of certain ORFs, while other ORFs with a less favorable codon composition may remain untranslated [53]. We have thus evaluated whether CUPrefs in *E5* resemble those in other *AlphaPV* genes. The *E5* genes exhibited CUPrefs similar to those in the early (*E6* and *E7*) genes (fig. 6), which are both implicated in oncogenesis. This is in line with previous work reporting that genes expressed at similar stages during viral infection have similar CUPrefs [25]. The observation that the *E5* ORFs are under purifying selection and the clustering of the CPUrefs of *E5* together with the two other oncogenes, reinforces the oncogenic role of the different E5 proteins in the life cycle of oncogenic human *AlphaPVs*.

In summary, our results strongly suggest that *E5* in *AlphaPVs* are *bona fide* genes and not merely spurious translations. This is supported by previous studies that already assigned different properties to *E5*, such as the alteration of membrane composition and dynamics [62, 60] and the down-regulation of surface MHC class I molecules [9, 10] for immune evasion. However, many questions about *E5* remain to be elucidated. Further experimental studies should be performed to provide evidence of the expression of the different *E5* ORFs *in vivo* and to elucidate whether *E5* originated through recombination, random nucleotide addition or another unknown mechanism.

## Data accessibility

Supplementary tables S1 and S2 are available online. The data from the Bali-Phy analyses and the random permutation test are available at https://github.com/anoukwillemsen/ONCOGENEVOL.

## Acknowledgements

This preprint has been reviewed and recommended by Peer Community In Evolutionary Biology (https://dx.doi.org/10.24072/pci.evolbiol.100067). We are grateful to the genotoul bioinformatics platform Toulouse Midi-Pyrenees (Bioinfo Genotoul) for providing computing and storage resources. The authors acknowledge the IRD itrop HPC (South Green Platform) at IRD montpellier for providing HPC resources that have contributed to the research results reported within this paper. This work was supported by the European Research Council Consolidator Grant CODOVIREVOL (Contract Number 647916) to IGB and by the European Union Horizon 2020 Marie Sklodowska-Curie research and innovation programme grant ONCOGENEVOL (Contract Number 750180) to AW.

## References

[1] Albà MM and Castresana J. Inverse relationship between evolutionary rate and age of mammalian genes. Molecular Biology and Evolution 22 (2005), 598–606. ISSN: 07374038. DOI: 10.1093/molbev/msi045.

[2] Ashby AD, Meagher L, Campo MS, and Finbow ME. E5 transforming proteins of papillomaviruses do not disturb the activity of the vacuolar H+-ATPase. Journal of General Virology 82 (2001), 2353–2362. ISSN: 00221317. DOI: 10.1099/0022-1317-82-10-2353.

[3] Ashrafi GH, Haghshenas MR, Marchetti B, O’Brien PM, and Campo MS. E5 protein of human papillomavirus type 16 selectively downregulates surface HLA class I. International Journal of Cancer 113 (2005), 276–283. ISSN: 0020-7136. DOI: 10.1002/ijc.20558.

[4] Ashrafi GH, Tsirimonaki E, Marchetti B, O’Brien PM, Sibbet GJ, Andrew L, and Campo MS. Down-regulation of MHC class I by bovine papillomavirus E5 oncoproteins. Oncogene 21 (2002), 248–259. ISSN: 0950-9232. DOI: 10.1038/sj.onc.1205008.

[5] Bennett MD, Woolford L, Stevens H, Van Ranst M, Oldfield T, Slaven M, O’Hara A J, Warren KS, and Nicholls PK. Genomic characterization of a novel virus found in papillomatous lesions from a southern brown bandicoot (Isoodon obesulus) in Western Australia. Virology 376 (2008), 173–182. ISSN: 00426822. DOI: 10.1016/j.virol.2008.03.014.

[6] Bravo IG and Alonso A. Mucosal Human Papillomaviruses Encode Four Different E5 Proteins Whose Chemistry and Phylogeny Correlate with Malignant or Benign Growth. Journal of Virology 78 (2004), 13613–13626. ISSN: 0022-538X. DOI: 10.1128/JVI.78.24.13613-13626.2004.

[7] Bravo IG and Felez-Sanchez M. Papillomaviruses: Viral evolution, cancer and evolutionary medicine. Evolution, Medicine and Public Health 2015 (2015), 32–51. ISSN: 20506201. DOI: 10.1093/emph/eov003.

[8] Cai J, Zhao R, Jiang H, and Wang W. De Novo Origination of a New Protein-Coding Gene in Saccharomyces cerevisiae. Genetics 179 (2008), 487–496. ISSN: 0016-6731. DOI: 10.1534/genetics.107.084491.

[9] Campo M, Graham S, Cortese M, Ashrafi G, Araibi E, Dornan E, Miners K, Nunes C, and Man S. HPV-16 E5 down-regulates expression of surface HLA class I and reduces recognition by CD8 T cells. Virology 407 (2010), 137–142. ISSN: 00426822. DOI: 10.1016/j. virol.2010.07.044.

[10] Cartin W and Alonso A. The human papillomavirus HPV2a E5 protein localizes to the Golgi apparatus and modulates signal transduction. Virology 314 (2003), 572–579. ISSN: 00426822. DOI: 10.1016/S0042-6822(03)00509-9.

[11] Carvunis A-R, Rolland T, Wapinski I, Calderwood MA, Yildirim MA, Simonis N, Charloteaux B, Hidalgo CA, Barbette J, Santhanam B, Brar GA, Weissman JS, Regev A, Thierry-Mieg N, Cusick ME, and Vidal M. Proto-genes and de novo gene birth. Nature 487 (2012), 370–374. ISSN: 0028-0836. DOI: 10.1038/nature11184.

[12] Castresana J. Selection of Conserved Blocks from Multiple Alignments for Their Use in Phylogenetic Analysis. Molecular Biology and Evolution 17 (2000), 540–552. ISSN: 0737-4038. DOI: 10.1093/oxfordjournals.molbev.a026334.

[13] Chacón KM, Petti LM, Scheideman EH, Pirazzoli V, Politi K, and DiMaio D. De novo selection of oncogenes. Proceedings of the National Academy of Sciences 111 (2014), E6–E14. ISSN: 0027-8424. DOI: 10.1073/pnas.1315298111.

[14] Chappell WH, Gautam D, Ok ST, Johnson BA, Anacker DC, and Moody CA. Homologous Recombination Repair Factors Rad51 and BRCA1 Are Necessary for Productive Replication of Human Papillomavirus 31. Journal of Virology 90 (2016). Ed. by Ross SR, 2639–2652. ISSN: 0022-538X. DOI: 10.1128/JVI.02495-15.

[15] Conrad M, Bubb VJ, and Schlegel R. The human papillomavirus type 6 and 16 E5 proteins are membrane-associated proteins which associate with the 16-kilodalton pore-forming protein. Journal of virology 67 (1993), 6170–8. ISSN: 0022-538X.

[16] Dasgupta S, Zabielski J, Simonsson M, and Burnett S. Rolling-circle replication of a high-copy BPV-1 plasmid. Journal of Molecular Biology 228 (1992), 1–6. ISSN: 00222836. DOI: 10.1016/0022-2836(92)90485-3.

[17] de Oliveira Martins L and Posada D. Infinitely long branches and an informal test of common ancestry. Biology Direct 11 (2016), 19. ISSN: 1745-6150. DOI: 10.1186/s13062-016-0120-y.

[18] de Oliveira Martins L and Posada D. Testing for Universal Common Ancestry. Systematic Biology 63 (2014), 838–842. ISSN: 1063-5157. DOI: 10.1093/sysbio/syu041.

[19] DiMaio D and Mattoon D. Mechanisms of cell transformation by papillomavirus E5 proteins. Oncogene 20 (2001), 7866–7873. ISSN: 0950-9232. DOI: 10.1038/sj.onc.1204915.

[20] DiMaio D and Petti LM. The E5 proteins. Virology 445 (2013), 99–114. ISSN: 00426822. DOI: 10.1016/j.virol.2013.05.006.

[21] Doorbar J, Quint W, Banks L, Bravo IG, Stoler M, Broker TR, and Stanley MA. The biology and life-cycle of human papillomaviruses. Vol. 30. SUPPL.5. 2012, F55–F70. ISBN: 1873-2518 (Electronic)\r0264-410X (Linking). DOI: 10.1016/j.vaccine.2012.06.083. arXiv: NIHMS150003.

[22] Doron-Faigenboim A and Pupko T. A Combined Empirical and Mechanistic Codon Model. Molecular Biology and Evolution 24 (2006), 388–397. ISSN: 0737-4038. DOI: 10.1093/molbev/msl175.

[23] Doron-Faigenboim A, Stern A, Mayrose I, Bacharach E, and Pupko T. Selection: A server for detecting evolutionary forces at a single amino-acid site. Bioinformatics 21 (2005), 2101–2103. ISSN: 13674803. DOI: 10.1093/bioinformatics/bti259.

[24] Edgar RC. MUSCLE: multiple sequence alignment with high accuracy and high throughput. Nucleic Acids Research 32 (2004), 1792–1797. ISSN: 1362-4962. DOI: 10.1093/nar/ gkh340.

[25] Félez-Sánchez M, Trösemeier J-H, Bedhomme S, González-Bravo MI, Kamp C, and Bravo IG. Cancer, Warts, or Asymptomatic Infections: Clinical Presentation Matches Codon Usage Preferences in Human Papillomaviruses. Genome Biology and Evolution 7 (2015), 2117–2135. ISSN: 1759-6653. DOI: 10.1093/gbe/evv129.

[26] Flores ER and Lambert PF. Evidence for a switch in the mode of human papillomavirus type 16 DNA replication during the viral life cycle. Journal of virology 71 (1997), 7167–79. ISSN: 0022-538X.

[27] Forman D, Martel C de, Lacey CJ, Soerjomataram I, Lortet-Tieulent J, Bruni L, Vignat J, Ferlay J, Bray F, Plummer M, and Franceschi S. Global Burden of Human Papillomavirus and Related Diseases. Vaccine 30 (2012), F12–F23. ISSN: 0264410X. DOI: 10.1016/j.vaccine. 2012.07.055.

[28] García-Pérez R, Ibáñez C, Godínez JM, Aréchiga N, Garin I, Pérez-Suárez G, Paz O de, Juste J, Echevarría JE, and Bravo IG. Novel Papillomaviruses in Free-Ranging Iberian Bats: No Virus–Host Co-evolution, No Strict Host Specificity, and Hints for Recombination. Genome Biology and Evolution 6 (2014), 94–104. ISSN: 1759-6653. DOI: 10.1093/gbe/evt211.

[29] Gillespie KA, Mehta KP, Laimins LA, and Moody CA. Human Papillomaviruses Recruit Cellular DNA Repair and Homologous Recombination Factors to Viral Replication Centers. Journal of Virology 86 (2012), 9520–9526. ISSN: 0022-538X. DOI: 10.1128/JVI.00247-12.

[30] Gottschling M, Bravo IG, Schulz E, Bracho MA, Deaville R, Jepson PD, Bressem M-FV, Stockfleth E, and Nindl I. Modular organizations of novel cetacean papillomaviruses. Molecular Phylogenetics and Evolution 59 (2011), 34–42. ISSN: 10557903. DOI: 10.1016/j. ympev.2010.12.013.

[31] Gottschling M, Göker M, Stamatakis A, Bininda-Emonds ORP, Nindl I, and Bravo IG. Quantifying the phylodynamic forces driving papillomavirus evolution. Molecular biology and evolution 28 (2011), 2101–13. ISSN: 1537-1719. DOI: 10.1093/molbev/msr030.

[32] Guindon S, Dufayard J-F, Lefort V, Anisimova M, Hordijk W, and Gascuel O. New Algorithms and Methods to Estimate Maximum-Likelihood Phylogenies: Assessing the Performance of PhyML 3.0. Systematic Biology 59 (2010), 307–321. ISSN: 1076-836X. DOI: 10.1093/sysbio/syq010.

[33] Hughes AL and Hughes MAK. Patterns of nucleotide difference in overlapping and non-overlapping reading frames of papillomavirus genomes. Virus research 113 (2005), 81–8. ISSN: 0168-1702. DOI: 10.1016/j.virusres.2005.03.030.

[34] Katoh K and Standley DM. MAFFT Multiple Sequence Alignment Software Version 7: Improvements in Performance and Usability. Molecular Biology and Evolution 30 (2013), 772–780. ISSN: 0737-4038. DOI: 10.1093/molbev/mst010.

[35] Koonin EV and Wolf YI. The common ancestry of life. Biology direct 5 (2010), 64. ISSN: 1745-6150. DOI: 10.1186/1745-6150-5-64.

[36] Kyte J and Doolittle RF. A simple method for displaying the hydropathic character of a protein. Journal of molecular biology 157 (1982), 105–32. ISSN: 0022-2836.

[37] McBride AA. Mechanisms and strategies of papillomavirus replication. Biological Chemistry 398 (2017), 919–927. ISSN: 14374315. DOI: 10.1515/hsz-2017-0113.

[38] Mehta K and Laimins L. Human Papillomaviruses Preferentially Recruit DNA Repair Factors to Viral Genomes for Rapid Repair and Amplification. mBio 9 (2018). Ed. by Imperiale MJ, e00064–18. ISSN: 2150-7511. DOI: 10.1128/mBio.00064-18.

[39] Moody CA and Laimins LA. Human papillomavirus oncoproteins: pathways to transformation. Nature Reviews Cancer 10 (2010), 550–560. ISSN: 1474-175X. DOI: 10.1038/ nrc2886.

[40] Münger K, Scheffner M, Huibregtse JM, and Howley PM. Interactions of HPV E6 and E7 oncoproteins with tumour suppressor gene products. Cancer surveys 12 (1992), 197–217. ISSN: 0261-2429.

[41] Narechania A, Chen Z, DeSalle R, and Burk RD. Phylogenetic incongruence among oncogenic genital alpha human papillomaviruses. Journal of virology 79 (2005), 15503–10. ISSN: 0022-538X. DOI: 10.1128/JVI.79.24.15503-15510.2005.

[42] Levine MT, Jones CD, Kern AD, Lindfors HA, and Begun DJ. Novel genes derived from noncoding DNA in Drosophila melanogaster are frequently X-linked and exhibit testis-biased expression. Proceedings of the National Academy of Sciences 103 (2006), 9935–9939. ISSN: 0027-8424. DOI: 10.1073/pnas.0509809103.

[43] Palmieri N, Kosiol C, and Schlötterer C. The life cycle of Drosophila orphan genes. eLife 3 (2014), e01311. ISSN: 2050-084X. DOI: 10.7554/eLife.01311.

[44] Petti LM, Reddy V, Smith SO, and DiMaio D. Identification of amino acids in the trans-membrane and juxtamembrane domains of the platelet-derived growth factor receptor required for productive interaction with the bovine papillomavirus E5 protein. Journal of virology 71 (1997), 7318–7327. ISSN: 0022-538X.

[45] Pim D, Collins M, and Banks L. Human papillomavirus type 16 E5 gene stimulates the transforming activity of the epidermal growth factor receptor. Oncogene 7 (1992), 27–32. ISSN: 0950-9232.

[46] R Core Team. R: A language and environment for statistical computing. Vienna, Austria, 2014.

[47] Rector A, Lemey P, Tachezy R, Mostmans S, Ghim S-J, Van Doorslaer K, Roelke M, Bush M, Montali RJ, Joslin J, Burk RD, Jenson AB, Sundberg JP, Shapiro B, and Van Ranst M. Ancient papillomavirus-host co-speciation in Felidae. Genome Biology 8 (2007), R57. ISSN: 14656906. DOI: 10.1186/gb-2007-8-4-r57.

[48] Rector A, Stevens H, Lacave G, Lemey P, Mostmans S, Salbany A, Vos M, Van Doorslaer K, Ghim S-J, Rehtanz M, Bossart GD, Jenson AB, and Van Ranst M. Genomic characterization of novel dolphin papillomaviruses provides indications for recombination within the Papillomaviridae. Virology 378 (2008), 151–161. ISSN: 00426822. DOI: 10.1016/j.virol. 2008.05.020.

[49] Redelings BD and Suchard MA. Joint bayesian estimation of alignment and phylogeny. Systematic Biology 54 (2005). Ed. by Lewis P, 401–418. ISSN: 10635157. DOI: 10.1080/ 10635150590947041.

[50] Robinson D and Foulds L. Comparison of phylogenetic trees. Mathematical Biosciences 53 (1981), 131–147. ISSN: 00255564. DOI: 10.1016/0025-5564(81)90043-2.

[51] Robles-Sikisaka R, Rivera R, Nollens HH, St. Leger J, Durden WN, Stolen M, Burchell J, and Wellehan JF. Evidence of recombination and positive selection in cetacean papillomaviruses. Virology 427 (2012), 189–197. ISSN: 00426822. DOI: 10.1016/j.virol.2012.01. 039.

[52] Roerink SF, Schendel R, and Tijsterman M. Polymerase theta-mediated end joining of replication-associated DNA breaks in C. elegans. Genome Research 24 (2014), 954–962. ISSN: 15495469. DOI: 10.1101/gr.170431.113.

[53] Ruiz-Orera J, Verdaguer-Grau P, Villanueva-Cañas JL, Messeguer X, and Albà MM. Translation of neutrally evolving peptides provides a basis for de novo gene evolution. 2 (2018), 1–7. ISSN: 2397334X. DOI: 10.1038/s41559-018-0506-6.

[54] Sakakibara N, Chen D, and McBride AA. Papillomaviruses Use Recombination-Dependent Replication to Vegetatively Amplify Their Genomes in Differentiated Cells. PLoS Pathogens 9 (2013), e1003321. ISSN: 1553-7374. DOI: 10.1371/journal.ppat.1003321.

[55] Schlötterer C. Genes from scratch–the evolutionary fate of de novo genes. Trends in genetics: TIG 31 (2015), 215–9. ISSN: 0168-9525. DOI: 10.1016/j.tig.2015.02.007.

[56] Schulz E, Gottschling M, Bravo IG, Wittstatt U, Stockfleth E, and Nindl I. Genomic characterization of the first insectivoran papillomavirus reveals an unusually long, second non-coding region and indicates a close relationship to Betapapillomavirus. Journal of General Virology 90 (2009), 626–633. ISSN: 0022-1317. DOI: 10.1099/vir.0.008011-0.

[57] Stamatakis A. RAxML version 8: a tool for phylogenetic analysis and post-analysis of large phylogenies. Bioinformatics 30 (2014), 1312–1313. ISSN: 1460-2059. DOI: 10.1093/ bioinformatics/btu033.

[58] Straight SW, Hinkle PM, Jewers RJ, and McCance DJ. The E5 oncoprotein of human papillomavirus type 16 transforms fibroblasts and effects the downregulation of the epidermal growth factor receptor in keratinocytes. Journal of virology 67 (1993), 4521–4532. ISSN: 0022-538X.

[59] Suchard MA and Redelings BD. BAli-Phy: simultaneous Bayesian inference of alignment and phylogeny. Bioinformatics 22 (2006), 2047–2048. ISSN: 1367-4803. DOI: 10.1093/ bioinformatics/btl175.

[60] Suprynowicz FA, Disbrow GL, Krawczyk E, Simic V, Lantzky K, and Schlegel R. HPV-16 E5 oncoprotein upregulates lipid raft components caveolin-1 and ganglioside GM1 at the plasma membrane of cervical cells. Oncogene 27 (2008), 1071–1078. ISSN: 0950-9232. DOI: 10.1038/sj.onc.1210725.

[61] Suyama M, Torrents D, and Bork P. PAL2NAL: robust conversion of protein sequence alignments into the corresponding codon alignments. Nucleic Acids Research 34 (2006), W609–W612. ISSN: 0305-1048. DOI: 10.1093/nar/gkl315.

[62] Bravo IG, Crusius K, and Alonso A. The E5 protein of the human papillomavirus type 16 modulates composition and dynamics of membrane lipids in keratinocytes. Archives of Virology 150 (2005), 231–246. ISSN: 0304-8608. DOI: 10.1007/s00705-004-0420-x.

[63] Theobald DL. On universal common ancestry, sequence similarity, and phylogenetic structure: the sins of P-values and the virtues of Bayesian evidence. Biology Direct 6 (2011), 60. ISSN: 1745-6150. DOI: 10.1186/1745-6150-6-60.

[64] Toll-Riera M, Bosch N, Bellora N, Castelo R, Armengol L, Estivill X, and Mar Albà M. Origin of primate orphan genes: A comparative genomics approach. Molecular Biology and Evolution 26 (2009), 603–612. ISSN: 07374038. DOI: 10.1093/molbev/msn281.

[65] Tomaić V. Functional Roles of E6 and E7 Oncoproteins in HPV-Induced Malignancies at Diverse Anatomical Sites. Cancers 8 (2016), 95. ISSN: 2072-6694. DOI: 10.3390/cancers8100095.

[66] Van Doorslaer K and McBride AA. Molecular archeological evidence in support of the repeated loss of a papillomavirus gene. Scientific Reports 6 (2016), 33028. ISSN: 2045-2322. DOI: 10.1038/srep33028.

[67] Venuti A, Paolini F, Nasir L, Corteggio A, Roperto S, Campo MS, and Borzacchiello G. Papillomavirus E5: The smallest oncoprotein with many functions. 10 (2011), 140. ISSN: 14764598. DOI: 10.1186/1476-4598-10-140.

[68] Wallace NA, Khanal S, Robinson KL, Wendel SO, Messer JJ, and Galloway DA. High-Risk Alphapapillomavirus Oncogenes Impair the Homologous Recombination Pathway. Journal of Virology 91 (2017). Ed. by Banks L, e01084–17. ISSN: 0022-538X. DOI: 10.1128/ JVI.01084-17.

[69] Wolf YI, Novichkov PS, Karev GP, Koonin EV, and Lipman DJ. The universal distribution of evolutionary rates of genes and distinct characteristics of eukaryotic genes of different apparent ages. Proceedings of the National Academy of Sciences 106 (2009), 7273–7280. ISSN: 0027-8424. DOI: 10.1073/pnas.0901808106.

[70] Woolford L, Rector A, Van Ranst M, Ducki A, Bennett MD, Nicholls PK, Warren KS, Swan RA, Wilcox GE, and O’Hara A J. A Novel Virus Detected in Papillomas and Carcinomas of the Endangered Western Barred Bandicoot (Perameles bougainville) Exhibits Genomic Features of both the Papillomaviridae and Polyomaviridae. Journal of Virology 81 (2007), 13280–13290. ISSN: 0022-538X. DOI: 10.1128/JVI.01662-07.

[71] Yang Z, Nielsen R, Goldman N, and Pedersen A-MK. Codon-Substitution Models for Heterogeneous Selection Pressure at Amino Acid Sites. Genetics 155 (2000), 431–49.

[72] Yonezawa T and Hasegawa M. Some problems in proving the existence of the universal common ancestor of life on Earth. TheScientificWorldJournal 2012 (2012), 479824. ISSN: 1537-744X. DOI: 10.1100/2012/479824.

[73] Yonezawa T and Hasegawa M. Was the universal common ancestry proved? Nature 468 (2010), E9–E9. ISSN: 0028-0836. DOI: 10.1038/nature09482.

[74] Zhou Q, Zhang G, Zhang Y, Xu S, Zhao R, Zhan Z, Li X, Ding Y, Yang S, and Wang W. On the origin of new genes in Drosophila. Genome Research 18 (2008), 1446–1455. ISSN: 1088-9051. DOI: 10.1101/gr.076588.108.

