## Supplementary figures and images for "Genome plasticity in Papillomaviruses and *de novo* emergence of *E5* oncogenes"

### fig. S1; fig. S2; fig. S3; fig. S4

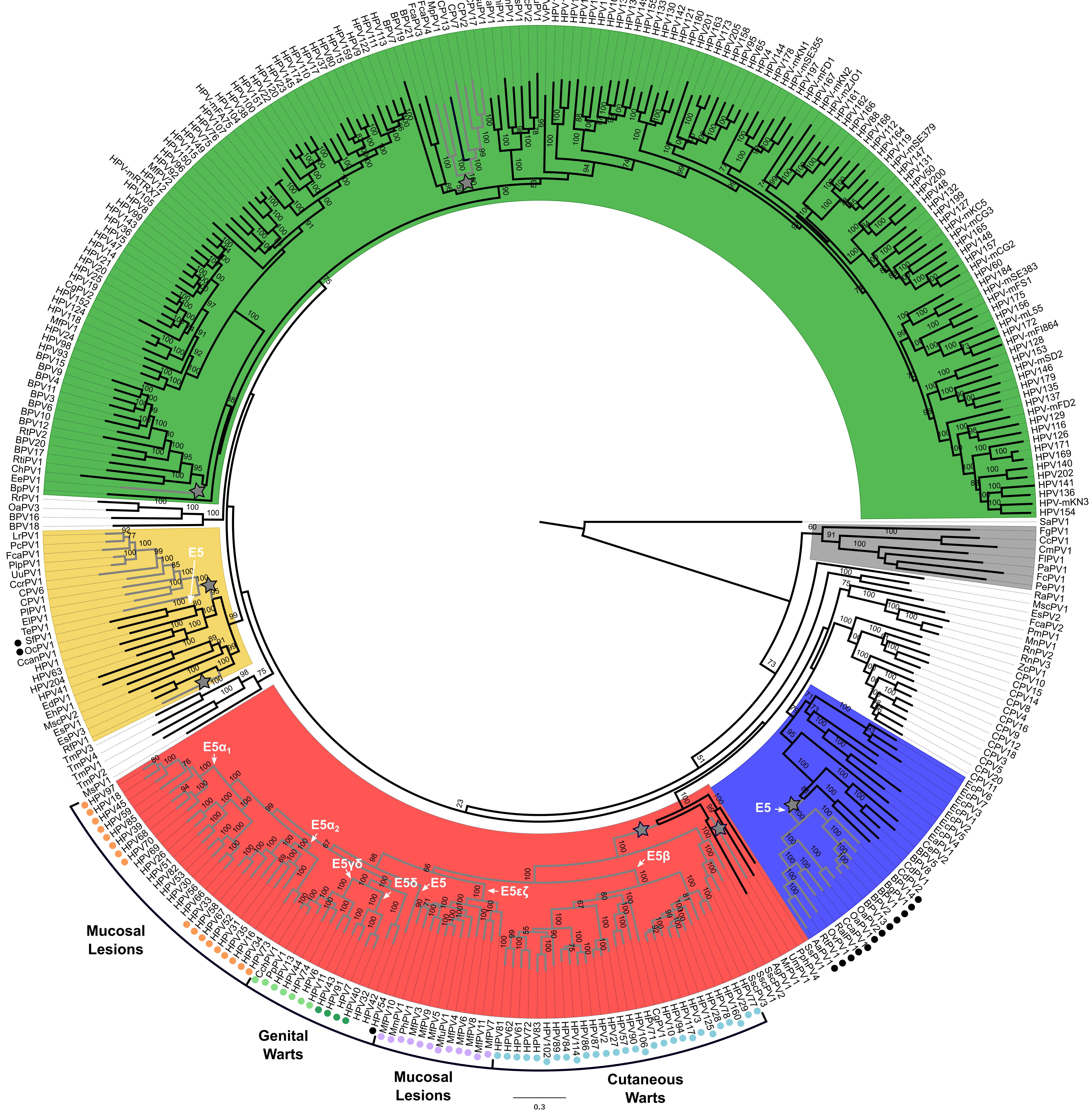

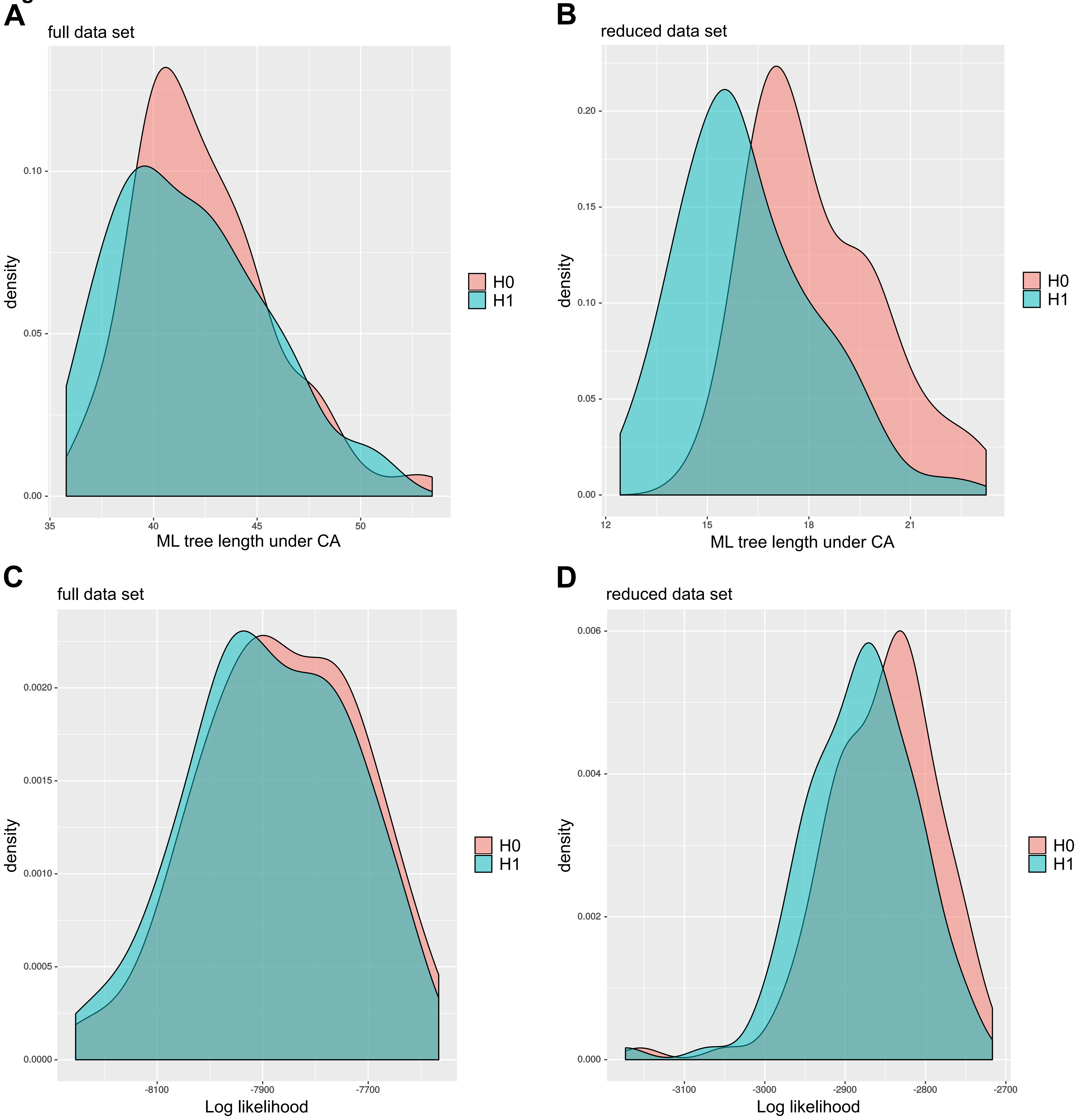

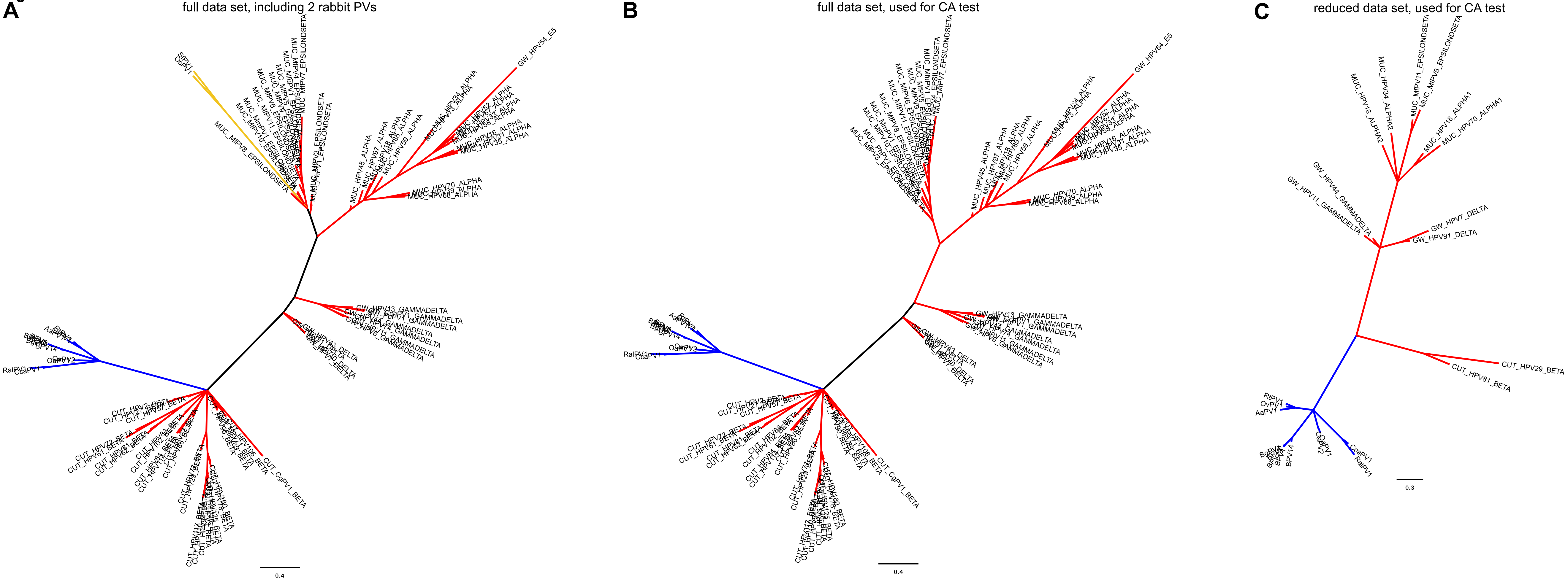

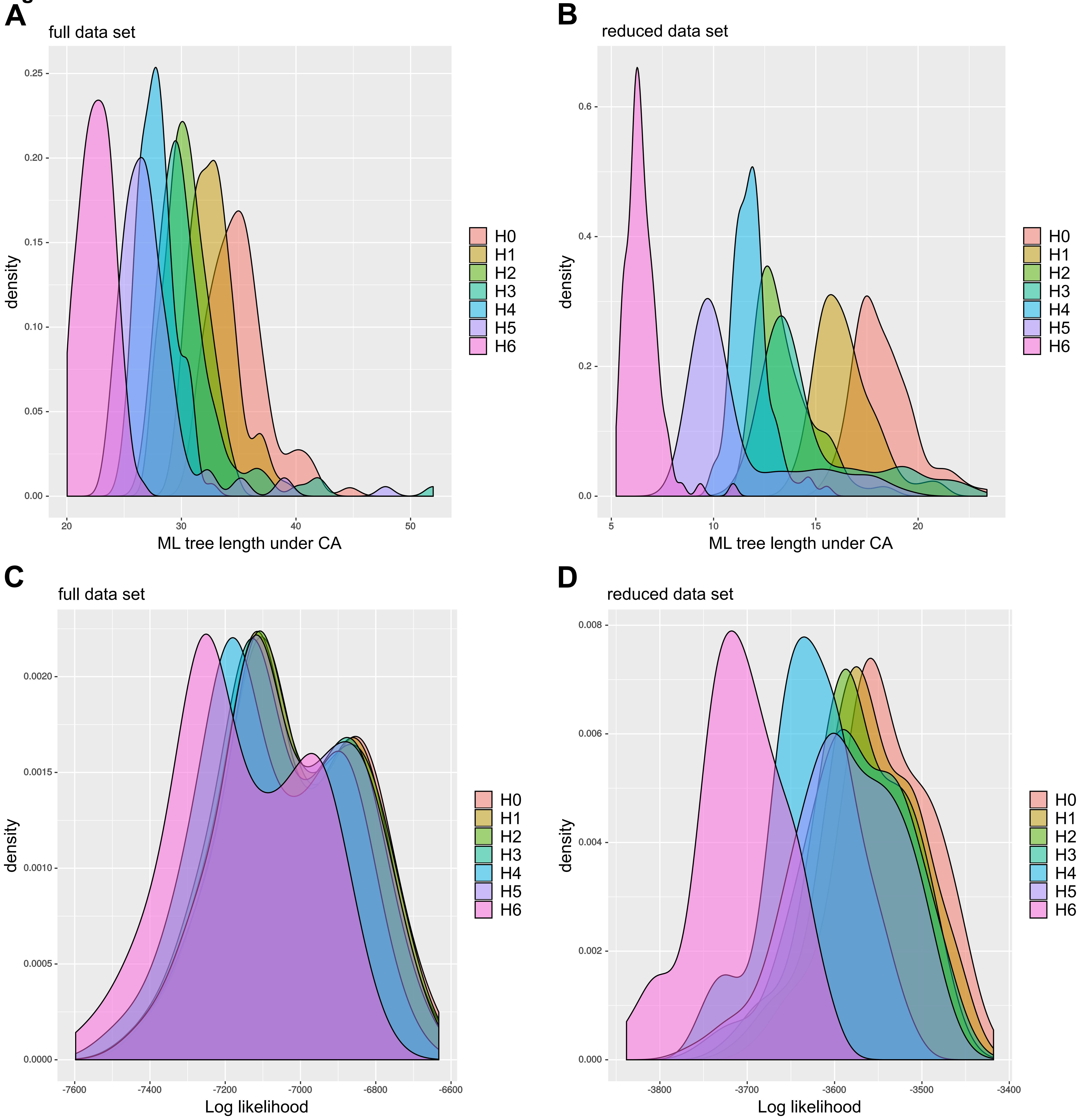
